# Improving the sensitivity of differential-expression analyses for under-powered RNA-seq experiments

**DOI:** 10.1101/2020.10.15.340737

**Authors:** Alex T. Kalinka

**Affiliations:** Milner Therapeutics Institute, Cambridge, UK

## Abstract

High-throughput studies, in which thousands of hypothesis tests are conducted simultaneously, can be under-powered when effect sizes are small and there are few replicates. Here, I describe an approach to estimate the FDR for a given experiment such that the ground truth is known. A decision boundary between true and false positive calls can then be learned from the data itself along the axes of fold change and expression level. By excluding hits that fall into the false positive space, the FDR of any given method can be controlled providing a means to employ less conservative methods for detecting differential expression without incurring the usual loss of precision. I show that coupling this approach with a feature-selection method - an elastic-net logistic regression - can increase sensitivity 10-fold above what is achievable with the prevailing methods of the day. An R package implementing these methods is available at https://github.com/alextkalinka/delboy.

## Introduction

Large-scale experiments are widely and routinely used in the biological and biomedical sciences and are arguably most often employed as exploratory tools. In this sense, they are used for ‘hypothesis generation’ and yet, as they have long been characterized as a simultaneous inference problem, the tools used to evaluate their output are individual-based ‘hypothesis tests’. This mismatch between experimental intent and statistical methodology has motivated attempts to define a more appropriate empirical null hypothesis (1).

For RNA-seq experiments, the problem is exacerbated by a majority of studies using three or fewer replicates (2). When the expected effect sizes are relatively small (e.g. median log fold change of 1 or less) then a given study can be severely under-powered from the individual hypothesis-test perspective. In these cases, the standard methods used to detect differentially expressed genes (DEGs), for example DESeq2 (3), will be conservative returning few or no hits after correcting for multiple testing. The value of the exploratory study, in this case, has been ablated as a result of the need to balance the sensitivity of a method with its false discovery rate (FDR).

If it were possible to predict which hits of a given method in conjunction with a given data-set were likely to be false positives, then we could tip the balance in favour of sensitivity without pushing the FDR unacceptably high. Here, I describe such a ‘cake-and-eat-it’ approach, the basic premise of which is that we are able to infer the FDR of a method using the input data itself, as opposed to any contrived data-set. By using the original data for evaluating the FDR, we retain the characteristics of the data that make the detection of true positives challenging, and ensure that our estimate of the FDR is as accurate as possible. I show that by coupling this empirical FDR estimation approach with a ‘feature-selection’ method (an elastic-net logistic regression) for discovering candidate hits, we are able to push the sensitivity for low-powered data 10-fold higher than established methods while keeping the FDR at acceptable levels.

## Methods

In the first section, I describe the empirical FDR estimation approach, broken down into its component steps. In the second section, I describe the elastic-net logistic regression for detecting candidate hits.

### Empirical FDR estimation

To be able to confidently infer the FDR using the original input data, we must control for the signal in the data that corresponds to the sample groupings. To do this, we apply a batch correction to the known experimental groups using an empirical Bayesian approach (4). This is a conservative approach, as the extent to which it fails to remove signal will only inflate the estimate of the FDR. This signal-corrected data is used as input when adding signal to the data to estimate the FDR.

However, before we can add known signal to a known set of genes, we first need to estimate both the number of non-null cases and the distribution of non-null log-fold changes in the original unmodified data. To do this, we use an empirical Bayesian approach to infer a mixture density of null and non-null cases in the original data (5). The input for estimating the number of non-null cases are the unadjusted p-values returned by DESeq2 run on the originsl data, and for estimating the distribution of log-fold changes, we use the log_2_ fold changes estimated by DESeq2.

Once we have the number of non-null cases (N) and their log-fold change distribution, we randomly, but reproducibly, sample N fold changes from the non-null distribution and attribute them to N randomly sampled genes. This signal is then added to the signal-corrected data using a binomial-thinning approach (6). This method adds signal to the data by sub-sampling counts using the binomial distribution.

With this ground-truth data in hand, we can now run any DEG-detection method on the data and determine its sensitivity and precision. In this case, we run both the elastic-net logistic regression hit-detection method (described below) and DESeq2 so that any inferiority relative to established methods can be reported back to the user.

Finally, we use the signal-controlled data to determine a decision boundary between true and false positive hits using a polynomial support vector machine (7) as implemented in the R package e1071. The two variables used for classification are the absolute log_2_ fold change and the log_10_ base mean expression level (per-gene), both taken from DESeq2 output for the signal-controlled data.

### Elastic-net logistic regression for hit detection

We employ a feature-selection approach to identify candidate hits, which makes the assumption that a reasonably large proportion of the genes are not differentially expressed. The method is the elastic-net penalized regression, so-called because it does not involve the stringency of penalty of a full lasso regression (keeping only one of a set of correlated predictors), but does not permit a large number of non-zero coefficients as does the ridge regression (8). A single parameter, α, determines the strength of the penalty with the lasso equal to 1 and the ridge equal to 0. In our case, we choose a value of 0.5.

We use the implementation in the R package glmnet and fit a logistic regression in which the response variable is the grouping of the data into one of at most two sample groups (e.g. control vs treatment) and the independent variable is the normalized expression of the gene - provided, for example, as transcripts per million (TPM), and, ideally quantified using a bias-aware RNA-seq quantification algorithm, such as kallisto (9) or salmon (10).

Once we have a set of hits, we apply the false positive classification learned from the signal-controlled data and annotate the hits as predicted to be true or false positives.

### RNA-seq data-set

To test the approach outlined above, we used an RNA-seq data-set in which, for a combination of biological and technical reasons, the fold changes were expected to be relatively small (11). The data was quantified into TPM values using salmon (10) and processed using delboy v0.0.1. A reproducible R markdown analysis script and all relevant data is available at https://github.com/alextkalinka/hormone-DE-analysis.

## Results

### FDR estimates and false positive classification

Performance estimates for delboy and DESeq2 are shown in Table 1. delboy exhibits a 10-fold higher sensitivity than DESeq2 but with a modest FDR of 4% after excluding hits predicted to be false positives.

**Table 1.** FDR estimates. Sensitivity and FDR estimates for delboy and DESeq2 when run on the same under-powered RNA-seq data-set with known signal added to a set of known genes. delboy estimates were taken after removing 6 hits predicted to be false positives. DESeq2 calls are all genes with adjusted p-values < 0.1. A total of 362 genes were determined to be non-null in the original data and had signal added to them using a binomial-thinning method.

| Algorithm | True calls | Sensitivity (%) | False calls | FDR (%) |
| --- | --- | --- | --- | --- |
| delboy | 72 | 20 | 3 | 4 |
| DESeq2 | 8 | 2 | 0 | 0 |

When run on the signal-controlled data, delboy’s false positives clustered together and had both low expression and low log-fold changes (Figure 1). Applying the false positive decision boundary to the original data, we find that 20 genes (out of 116 in total) fall into the false positive space.

**Figure 1.**
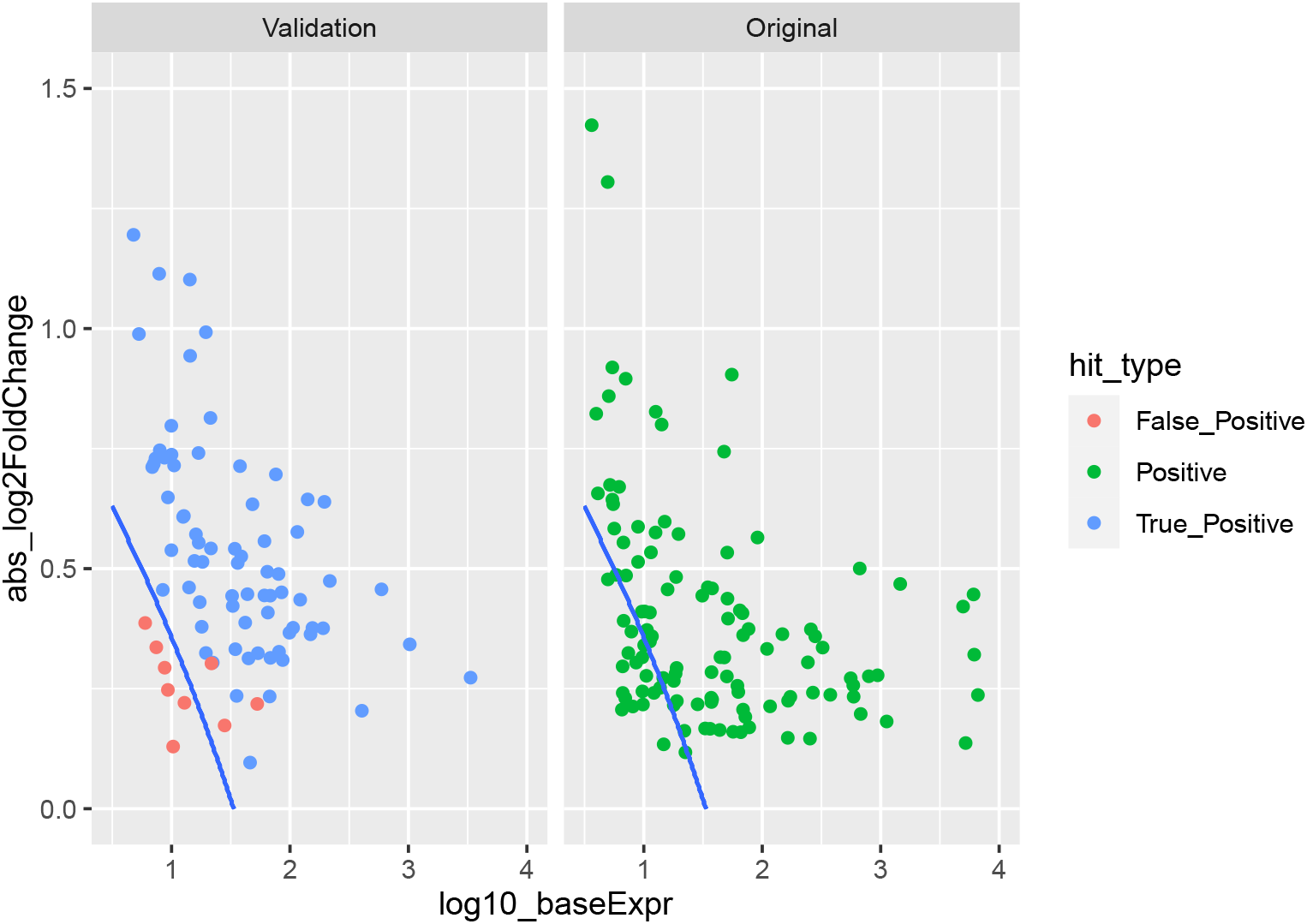
False positive classification using delboy. The panel on the left shows signal-corrected ‘validation’ data, and the panel on the right shows the original, unmodified data. A polynomial support vector machine decision boundary is shown with predicted false positives falling below it. For the original data, 20 genes fall into the false positive space.

It is worth noting that delboy’s false negatives fall predominantly in the false positive space predicted by the SVM (Figure 2). Hence, they were likely not called due to having low expression and low fold changes. This suggests that we can think of the false positive boundary as a limit of detection, one which is a function of both expression level and fold change.

**Figure 2.**
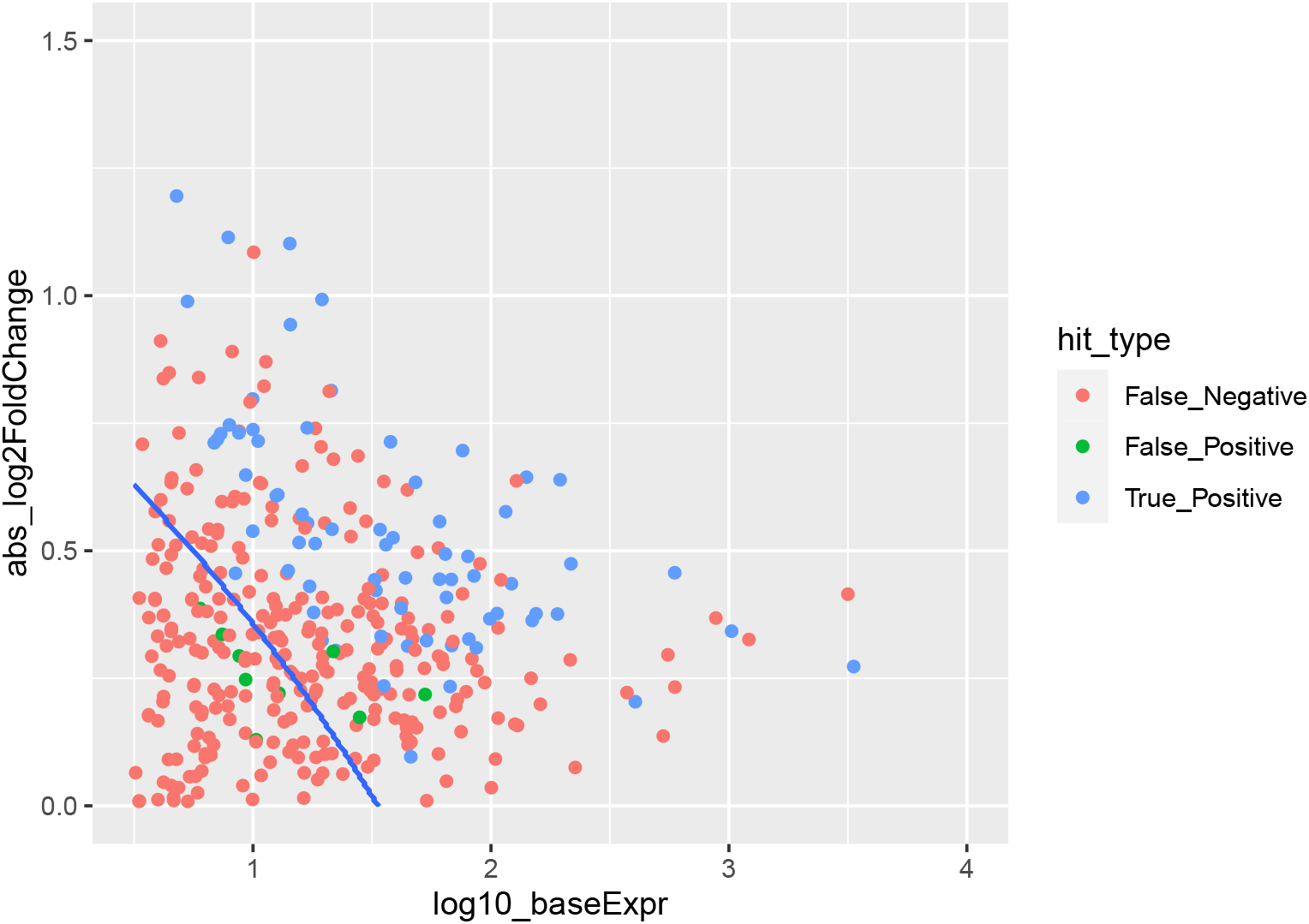
False negatives in relation to false positive boundary. For the signal-controlled data, false negatives fall predominantly below the false positive decision boundary.

## Conclusions

Using an empirical estimate of the FDR of an RNA-seq experiment using the data itself allows for false positives to be characterised in terms of fold changes and expression level. By excluding predicted false positives, we are empowered to use less conservative DEG-detection methods that boost our sensitivity while keeping the FDR low.

## Supporting Information

The delboy algorithm R package is available at https://github.com/alextkalinka/delboy. A reproducible analysis script and relevant data is available at https://github.com/alextkalinka/hormone-DE-analysis.

## Acknowledgments

Thanks are due to Iva Kelava and Madeline Lancaster for access to unpublished data.

## Notes

https://github.com/alextkalinka/delboy

### Competing Interest Statement

The authors have declared no competing interest.

## References

1. Efron B (2004) Large-scale simultaneous hypothesis testing. Journal of the American Statistical Association 99(465):96–104.

2. Li Y, et al. (2020) Decode-seq: a practical approach to improve differential gene expression analysis. Genome Biology 21(1).

3. Love MI, Huber W, Anders S (2014) Moderated estimation of fold change and dispersion for RNA-seq data with DESeq2. Genome Biology 15(12).

4. Johnson WE, Li C, Rabinovic A (2006) Adjusting batch effects in microarray expression data using empirical bayes methods. Biostatistics 8(1):118–127.

5. Efron B (2007) Size, power and false discovery rates. The Annals of Statistics 35(4):1351–1377.

6. Gerard D (2020) Data-based RNA-seq simulations by binomial thinning. BMC Bioinformatics 21(1).

7. Chang CC, Lin CJ (2011) LIBSVM. ACM Transactions on Intelligent Systems and Technology 2(3):1–27.

8. Friedman J, Hastie T, Tibshirani R (2010) Regularization paths for generalized linear models via coordinate descent. Journal of Statistical Software 33(1).

9. Bray NL, Pimentel H, Melsted P, Pachter L (2016) Near-optimal probabilistic RNA-seq quantification. Nature Biotechnology 34(5):525–527.

10. Patro R, Duggal G, Love MI, Irizarry RA, Kingsford C (2017) Salmon provides fast and bias-aware quantification of transcript expression. Nature Methods 14(4):417–419.

11. Kelava I, Chiaradia I, Pellegrini L, Kalinka AT, Lancaster M (2020) Sex hormones influence excitatory neuron production during human brain development. bioRxiv.

